# Single-cell *in vivo* imaging reveals light-initiated circadian oscillators in zebrafish

**DOI:** 10.1101/716100

**Authors:** Haifang Wang, Zeyong Yang, Xingxing Li, Dengfeng Huang, Shuguang Yu, Jie He, Yuanhai Li, Jun Yan

## Abstract

Circadian clock is a cell-autonomous time-keeping mechanism established gradually during embryonic development. Here we generated a transgenic zebrafish line carrying a destabilized fluorescent protein driven by the promoter of a core clock gene, *nr1d1*, to report *in vivo* circadian rhythm at the single-cell level. By time-lapse imaging of this fish line, we observed the sequential initiation of the reporter expression starting at photoreceptors in pineal gland then spreading to cells in other brain regions. Even within pineal gland, we found heterogeneous onset of *nr1d1* expression in which each cell undergoes circadian oscillation superimposed over cell-type specific developmental trajectory. Furthermore, we found that single-cell expression of *nr1d1* showed synchronous circadian oscillation under light-dark cycle. Remarkably, single-cell oscillations were lost in animals raised under constant darkness while developmental trend still persists. It suggests that light exposure in early clock development initializes cellular clocks rather than synchronizes existing individual oscillators as previously believed.

## Introduction

Circadian rhythm evolves to align animal behaviors to periodic daily environmental changes. At the molecular level, vertebrate circadian clock is mainly generated through transcriptional/translational feedback loops of core clock genes [1]. Among them, two transcription factors (TFs), BMAL1 (also known as ARNTL or MOP3) and CLOCK form heterodimers to bind to E-boxes in the promoters and initiate the transcription of their target genes [2–4] including Per family genes (Per1, Per2, and Per3) and Cry family genes (Cry1 and Cry2). The activation of these genes results in the formation of PER/CRY complex and thereby inhibits CLOCK/BMAL1 transcriptional activity, forming a negative feedback loop [5]. Nuclear receptor, REV-ERB α (also known as NR1D1), represses the transcription of Bmal1 and itself is under the transcriptional regulation of BMAL1/CLOCK giving rise to second negative feedback loop of circadian clock. The genome-wide regulation by circadian TFs such as BMAL1/CLOCK and REV-ERB α typically leads to thousands of genes showing circadian expression in a given tissue. Although the basic network of core circadian genes is present in almost every cell, many of the circadian-controlled genes are tissue-specific or cell-type specific. Their circadian expression is a result of either tissue-specific binding of circadian TFs [6] or transcriptional cascade from tissue-specific TFs regulated by circadian TFs [7]. At the organismal level, the overt circadian rhythm is governed by an intricate network of circadian oscillators, in which the master pacemaker such as SCN in mammals or pineal gland in zebrafish is believed to play a pivotal role [1,8]. Traditional *in vivo* or *ex vivo* studies of cellular circadian clock have relied on luciferase reporter systems driven by core clock gene promoters [9,10]. But luciferase imaging can only achieve single-cell resolution in organotypic slices in culture with high-resolution CCD camera. Transgenic zebrafish lines carrying luciferase reporters driven by core clock gene promoters have been developed in larval zebrafish. But *in vivo* luciferase imaging of these fish lines lacked single-cell resolution and can only report the population-level circadian rhythm [9,11]. Genetically encoded calcium indicator (GCaMP) has been widely used to monitor *in vivo* single-cell calcium activity. In comparison, there is still a lack of zebrafish line to report the circadian expression at the single cell level *in vivo*.

Circadian rhythm has to be established at molecular, cellular, tissue and behavioral levels during animal development. Day-night rhythms in the fetal rat SCN are first detected between E19 and E21 [12]. Rhythmic expression of circadian clock genes is not detected in both *in vitro* and *in vivo* mouse embryonic stem cells (ESCs) but only appears when ESCs differentiate into neural stem cells [13,14]. By *ex vivo* luciferase imaging of mouse fetal SCN, Carmona-Alcocer et al. showed that a few cells in SCN start circadian oscillations on E14.5, widespread synchronized oscillations were formed on E15.5, and then dorsal-ventral phase wave was established at P2 [15]. Zebrafish (*Danio rerio*) embryos that are *in vitro* fertilized and transparent provide an accessible model organism for *in vivo* live imaging, and thereby being widely used in the study of animal development. A functional circadian clock, characterized by free-running activity, rhythmic cell cycle and circadian gene expression, is established after hatching in zebrafish [16]. However, how single-cell circadian clocks were established during embryonic development is still unclear. It is well-documented that early exposure to light-dark (LD) cycle is required for the development of circadian clock in zebrafish larva [9]. But it is still under debate whether the effect of light on clock development is a synchronization of already existing oscillators or an initiation of single-cell clocks. Questions like these can only be addressed by *in vivo* imaging of single-cell circadian clock reporter.

Here we report a transgenic zebrafish line using destabilized fluorescent protein, Venus-NLS-PEST (VNP), driven by the promoter of a key circadian clock gene, *nr1d1*. This system allows us to monitor the development of single-cell circadian rhythm in live zebrafish larva in a cell-type specific manner. We observed that VNP reporter expression undergoes stepwise onset starting at the photoreceptor cells in pineal gland then spread to cells in other brain regions. Using single-cell RNAseq, we characterized the cell types expressing VNP in the whole brain. Within pineal gland, we found that each cell undergoes circadian oscillation superimposed over cell-type specific developmental trajectories. Under LD cycle, cellular expression of *VNP* shows synchronous circadian oscillation. However, the circadian expression of *nr1d1* positive cells was lost while the developmental trend was still present at single cell level in fish raised under constant darkness. Our result suggests that the effect of early exposure of LD cycle on clock ontogeny is an initiation of functional single-cell oscillators rather than a synchronization of already existing oscillators.

## Results

### Screening for *in vivo* circadian reporters in zebrafish

To monitor circadian rhythm at the single-cell level in live animals, we have screened for *in vivo* circadian reporters among various combinations of destabilized fluorescent proteins driven by core clock gene promoters in larval zebrafish. We first tested peTurboGFP-dest1 (TGFPD1) encoding a destabilized variant of green fluorescent protein TurboGFP. The plasmids of *bmal1a/bmal2/per2/nr1d1*:TGFPD1 were constructed by homologous recombination. We observed that none of these plasmids were expressed in zebrafish embryos. We then tested the plasmids containing core clock gene promoters driven Venus-NLS-PEST (VNP), another form of destabilized fluorescent protein. We found that the plasmids of *cry2b/bmal1a/per2*:VNP were not expressed in zebrafish embryos, while *per1a*:*VNP* has too low expression to be used for *in vivo* imaging. For *nr1d1*:VNP, it has been reported that zebrafish *nr1d1* gene has two promoters, i.e. ZfP1 (distal) and ZfP2 (proximal). ZfP1 is conserved and functionally similar to mammalian *Nr1d1* promoter [17]. We found that the fish line containing only the proximal *nr1d1* promoter ZfP2 (1.5kb) failed to drive VNP expression. But *nr1d1*:VNP containing both ZfP1 and ZfP2 (6.2kb) showed robust expression in zebrafish embryos (Fig. 1a and Supplementary Table 1). Therefore, we chose *nr1d1*:VNP containing ZfP1 and ZfP2 promoters as the circadian reporter and generated the transgenic fish line Tg(*nr1d1*:VNP) by the Tol2 system.

**Fig. 1.**
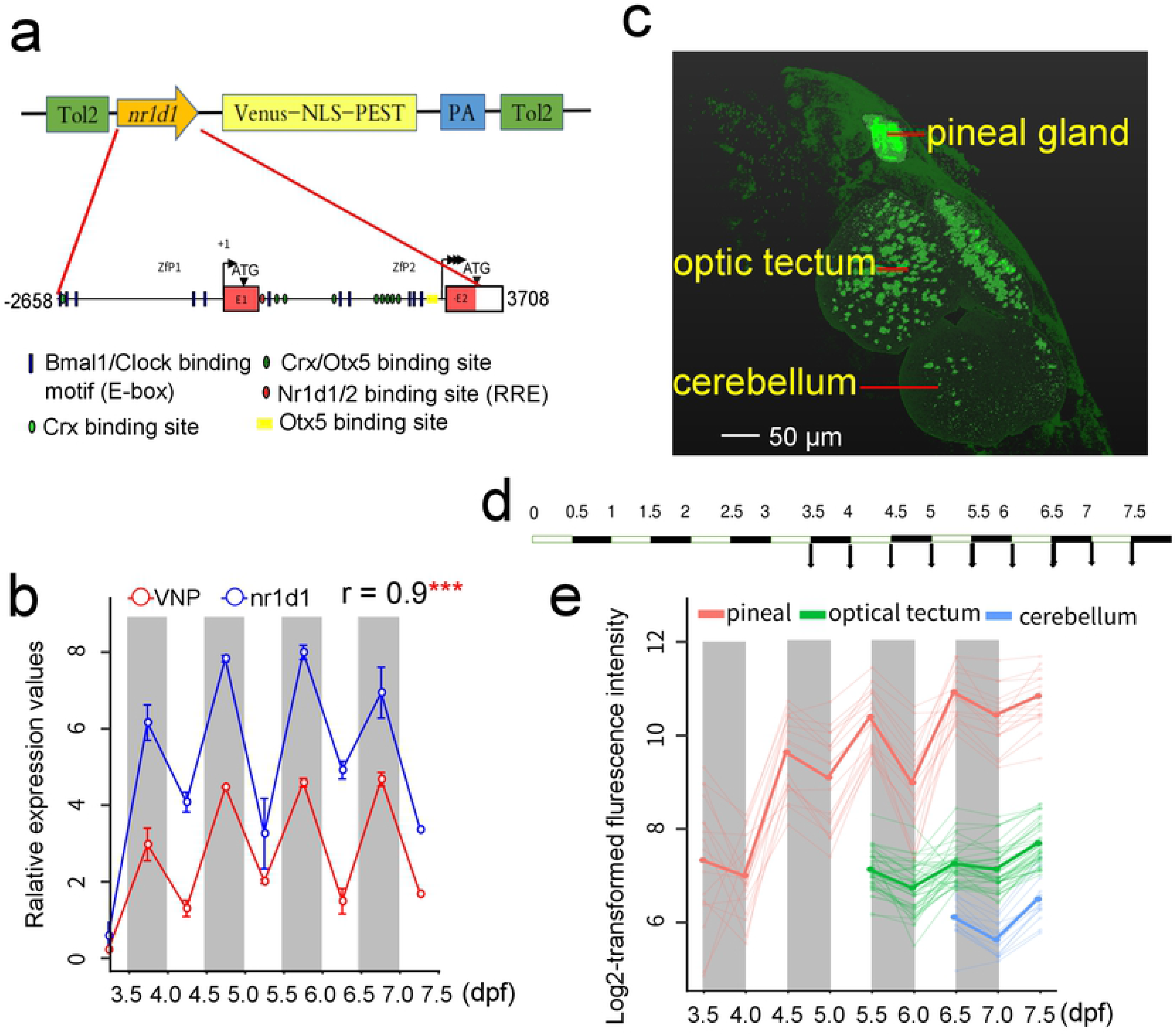
Construction of *in vivo* circadian reporter in zebrafish. (a) The upper graph shows the schematic of *nr1d1*:VNP construct design. The lower graph shows a magnified view of the *nr1d1* promoter sequence used for driving the circadian expression of VNP. The putative RRE (Nr1d1/2 binding site), E-box (Bmal1/Clock binding site), Crx, Otx5 and Crx/Otx5 binding sites were indicated in blue rectangle, red oval, green oval, yellow rectangle, and dark green oval respectively. (b) Plot of real-time PCR results of *nr1d1* and *Venus*-*pest* expression. Each time point has 2 replicas, and each replicate is the pool of 7-10 fish. The dots show the original value, while the solid lines and the bars represent mean ± sem. (c) Representative image shows the spatial distribution of *nr1d1*:VNP-positive cells in different brain regions at 7.5dpf. (d) Experimental design to examine the developmental dynamics of *nr1d1*:VNP expression in the whole zebrafish brain. (e) Single-cell tracing result of *nr1d1*:VNP-positive cells in the pineal gland (red), the optic tectum (green), and the cerebellum (blue) for one fish. The cells were tracked from 3.5dpf in pineal gland, 5.5dpf in optic tectum and 6.5dpf in cerebellum respectively. Each thin line represents one cell while the thick line represents the mean value of cells in each brain region.

The 6.2-kb ZfP1 and ZfP2 promoter region to drive VNP expression includes the entire set of known *cis*-regulatory elements: E-box, RRE, Crx- and Otx5-binding sites (Fig. 1a). We measured the mRNA levels of endogenous *nr1d1* and VNP expression in the Tg(*nr1d1*:VNP) transgenic fish from 3.5 dpf (days post-fertilization) to 7.5 dpf under the 12h/12h LD cycles using real-time PCR. *nr1d1* and VNP showed highly correlated expression patterns (Pearson’s r = 0.9) indicating that VNP expression faithfully reported the expression of *nr1d1* at the mRNA level. In addition, both genes showed higher expression level at the dawn than dusk over days (Fig. 1b), which is consistent with our previous result that *nr1d1* shows circadian expression peaking at ZT23 (Zeitgeber time) in larval zebrafish [7]. We examined the spatial distribution of fluorescence-labeled cells in *nr1d1*:VNP at 7.5 dpf using *in vivo* two-photon imaging system and found *nr1d1*:VNP-positive cells in many brain regions including the pineal gland, the optic tectum, and the cerebellum (Fig. 1c, Supplementary Movie 1,2).

We next monitored *in vivo* expression of *nr1d1*:VNP at the whole-brain scale from 3.5 dpf to 7.5 dpf at ZT0 and ZT12 (Fig. 1d) by time lapse imaging. The expression of *nr1d1*:VNP was sequentially turned on and underwent gradual increase in distinct brain regions during development, as early as 3.5 dpf in the pineal gland, followed by the optic tectum at 5.5 dpf, and other brain regions such as the cerebellum at later time points (Fig. 1e Supplementary Table 2, Supplementary Movie 1,2). In addition, single-cell tracing revealed that circadian oscillations of *nr1d1*:VNP expression appear to peak at ZT12 across brain regions (Fig. 1e). The expression pattern of *nr1d1*:VNP closely resembled that of endogenous *nr1d1* reported by Delaunay et al. [18]. So *nr1d1*:VNP fish can be used as a reporter for both developmental expression and circadian expression of endogenous *nr1d1* gene in zebrafish.

### Characterization of *nr1d1*:VNP expressing cells

To identify the cell types expressing *nr1d1*:VNP in the whole brain, we conducted single cell RNA-seq (scRNA-seq) of ~15,000 cells dissociated from the brain of Tg(*nr1d1*:VNP) larval fish at 6.5dpf. Among them, 6514 cells were identified with more than 500 genes and were used for the downstream analysis. In total, 26 cells clusters were classified from scRNAseq. They were manually annotated by comparing the marker genes with those from whole-brain single-cell RNA-seq data in adult zebrafish [19] (Fig. 2a). We found that the mRNA of *nr1d1*:VNP was enriched (adjust p-value < 0.01) in cell clusters of photoreceptors in pineal gland, glutamate neurons in forebrain and hypothalamus, granule cells in cerebellum, dorsal habenula cells as well as some non-neuron cells such as oligodendrocyte, retinal-pigment epithelium-like cell, epidemis, and endothelial cells (Fig. 2b).

**Fig. 2.**
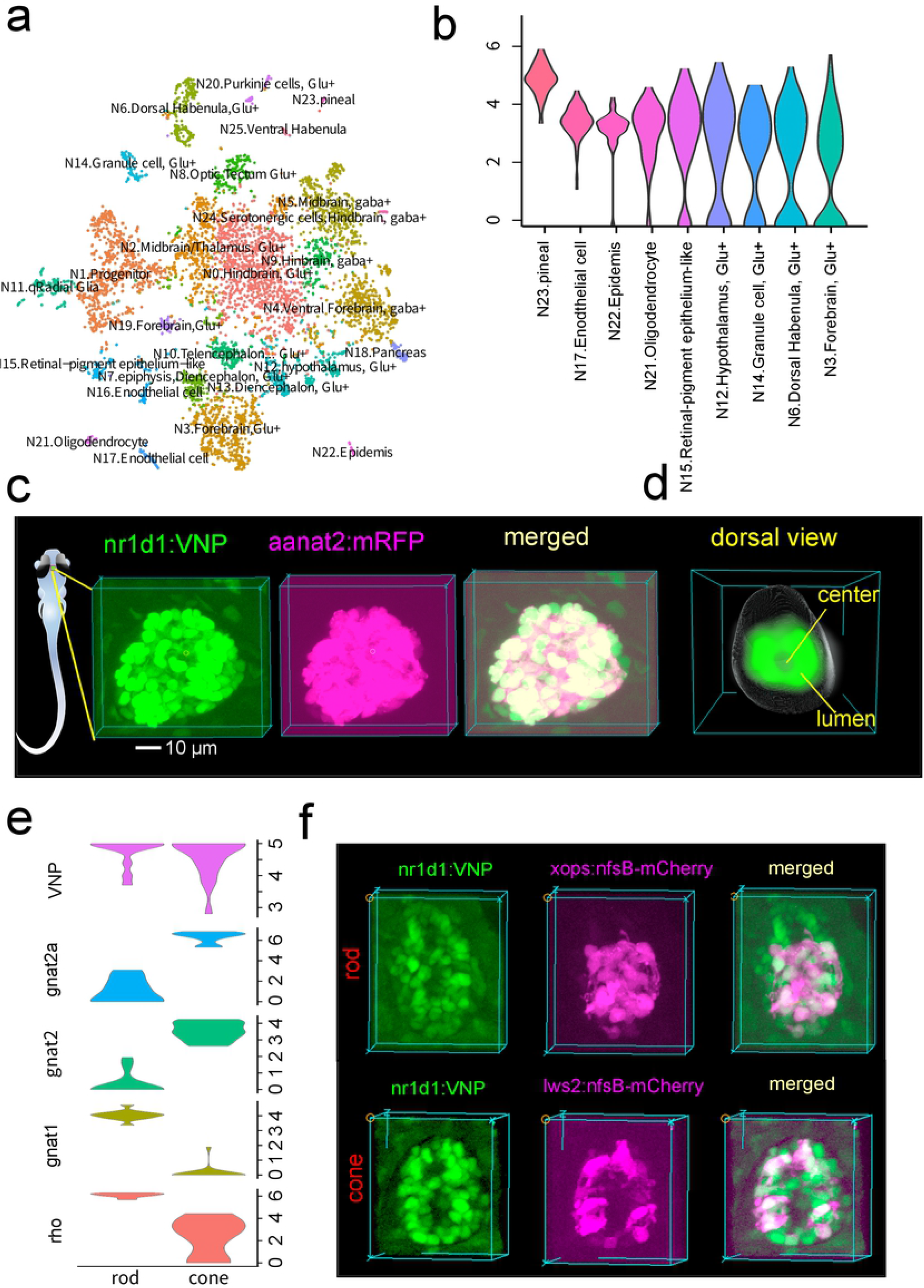
Characterization of *nr1d1*:VNP expressing cells. **(a)** t-SNE visualization of single cell clusters. The clusters were annotated by comparing to the adult zebrafish single cell RNA-seq data. (b) Violin plot demostrated the clusters with enriched expression of *nr1d1*:VNP. (c) Fluorescence images show the co-expression of *nr1d1*:VNP (nuclear signal) and *aanat2*:mRFP (cytoplasmic signal) in the zebrafish pineal gland. The left graph illustrates the 3D location of pineal gland in zebrafish larva. (d) 3D reconstruction of a common zebrafish pineal gland by aligning and averaging of six fish. The grey sphere represents the boundary of pineal gland and the green color represents the density distribution of *nr1d1*:VNP-positive cells in the pineal gland. (e) Violin plot showed the expression of rod cell markers (*rho*, *gnat1*), cone markers (*gnat2*, *gnat2a*), and *nr1d1*:VNP in photoreceptor clusters. (e) Fluorescence images showed the co-expression of *nr1d1*:VNP (nuclear signal) with rod fish line Tg(*xops*:nfsB-mCherry) (cytoplasmic signal) and cone fish line Tg(*lws2*:nfsB-mCherry) (cytoplasmic signal) in the zebrafish pineal gland.

We next examined the expression of *nr1d1*:VNP in pineal gland at the single-cell resolution. Fig. 2c showed an example of 3D reconstruction of *nr1d1*:VNP signals in the pineal gland. At 4.5 dpf, each pineal gland contains about 60 *nr1d1*:VNP-positive cells. To better understand the cell type of these cells, we crossed Tg(*nr1d1*:VNP) with Tg(*aanat2*:mRFP) [11]. Averaged 3D cell density distribution of *nr1d1*:VNP-positive cells across six fish (Methods section) revealed that *nr1d1*:VNP-positive cells were mostly distributed around the lumen of the pineal gland while the cell density was relatively low in the central part of the pineal gland (Fig. 2d). We observed that all *nr1d1*:VNP signals were overlapped with the *aanat2*:mRFP signals indicating that all *nr1d1*:VNP-positive cells were melatonin-synthesizing photoreceptor cells in the pineal gland (Fig. 2c, Supplementary Movie 3). According to our scRNA-seq data, VNP is expressed in both rod photoreceptors and cone photoreceptors (Fig 2e). Indeed, we observed co-labeling of VNP with rod cell marker (*xops*) and cone cell marker (*lws2*) in pineal gland by crossing Tg(*nr1d1*:VNP) fish line with rod fish line Tg(*xops*:nfsB-mCherry) and cone fish line Tg(*lws2*:nfsB-mCherry), respectively (Fig 2f). Our scRNA-seq data also suggested that *nr1d1*:VNP is expressed in proliferative cells. Indeed, when we crossed our VNP fish line with Tg(*her4*:DsRed) zebrafish marking proliferative cells [20], we found co-labeling of VNP and DsRed (Supplementary Fig.1).

### Developmental dynamics of *nr1d1*:VNP expression within pineal gland

To examine the development of circadian oscillations more closely in pineal gland, we imaged *nr1d1*:VNP signals in the pineal gland from 3.5 dpf to 6.5 dpf using two-photon imaging (Fig. 3a, b, Supplementary Movie 4-9, Supplementary Table 5). The majority of cells already showed higher *nr1d1*:VNP signals in dusk than dawn when the signals first become detectable in the pineal gland (Supplementary Fig. 2). This pattern does not change significantly during development (ANOVA P = 0.78) (Supplementary Fig. 2). However, we found that VNP positive cells within pineal gland display heterogeneous temporal patterns of *nr1d1*:VNP expression over development (Fig. 3c). Some cells showed rapid increase in the baseline level while other cells showed more robust circadian oscillations. We then quantified the circadian and developmental components in each cell by fitting the time-series data with a regression model that combines a step-wise function of time, (B*(−1)^(x+1)^) denoting circadian oscillation, with a linear function of time, (Ax + C) denoting the developmental increase (Fig. 3d). After fitting the model, we found that the regression coefficients of developmental effect (A) and circadian oscillation (B) both vary widely across cells but there is a negative correlation between them (Fig. 3e).

**Fig. 3.**
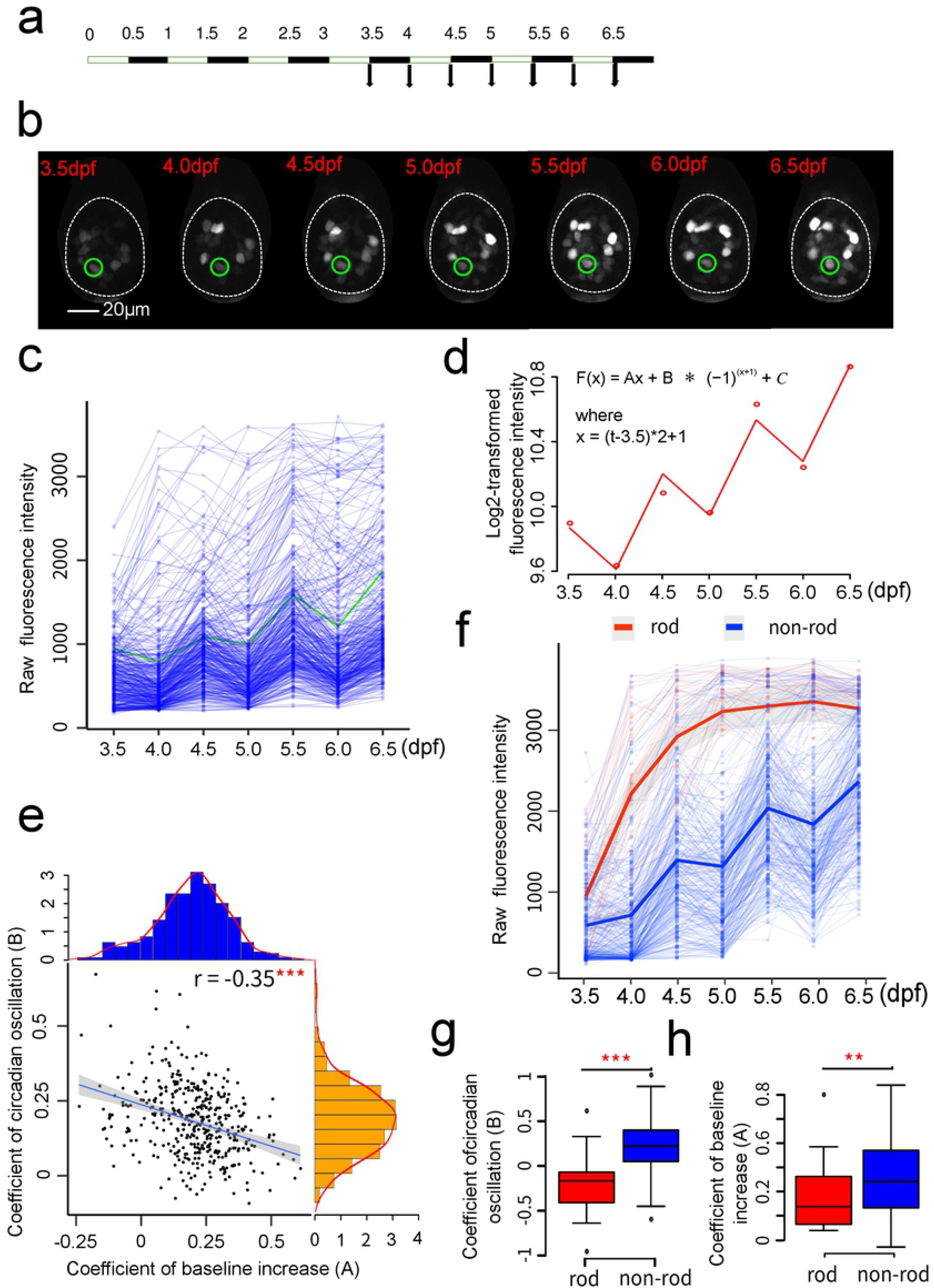
Developmental dynamics of *nr1d1*:VNP expression. (a) Experimental design to examine the developmental dynamics of *nr1d1*:VNP expression. (b) Fluorescence images illustrate the tracing of one *nr1d1*:VNP cell during development from 3.5 dpf to 6.5 dpf. (c) Raw fluorescence intensity of the traced cells. Representative cell in (b) was highlighted in green (six fish). (d) Regression model used to fit the fluorescence intensity. The dots represent the fluorescence intensity, while the line represent the fitting curve. (e) Scatter plot demonstrated the relationship between coefficients of developmental effect (A) and circadian oscillation (B). The histograms in blue and orange showed the distribution of coefficients of developmental effect (A) and the distribution of coefficients of circadian oscillations (B), respectively. *** represents Pearson’s correlation p-value <0.001. (f) Raw fluorescence intensity of rod cells and non-rod cells (six fish). Each thin line represents one cell and each dot represents raw fluorescence intensity. Thick lines represent the loess-smoothed curvess for all rod cells in red and non-rod cells in blue respectively. The shaded areas show the 95% confidence level of the smoothed curve. (g) A comparison of circadian oscillation coefficient (B) between rod and non-rod cells. Two-tailed Student’s t-test was applied to calculate the levels of significance between the two types of cells. *** represents P<0.001. (g) A comparison of developmental coefficient (A) between rod and non-rod cells. Two-tailed Student’s t-test was applied to calculate the levels of significance between the two types of cells. ** represents P<0.01.

To reveal if the heterogeneity of single-cell temporal profiles within pineal gland are due to differences in cell type, we next conducted cell-type specific imaging by crossing *nr1d1*:VNP with the fish line labeling rod cells in red fluorescent protein Tg(*xops*:nfsB-mCherry). After imaging *nr1d1*:VNP signals in the rod and non-rod cells simultaneously in the same fish during development from 3.5 dpf to 6.5 dpf (Supplementary Table 9, Supplementary Movie 25-30), we found that rod cells have a faster rate of developmental increase in *nr1d1*:VNP signals (A) but lower amplitude of circadian oscillation (B) than non-rod cells (Fig. 3f-h). Taken together, pineal photoreceptor cells of zebrafish larvae undergo circadian oscillations superimposed over cell-type specific developmental trajectories.

### Single-cell circadian oscillations in pineal gland

We next monitored the *nr1d1*:VNP signals in the pineal glands of live zebrafish larvae with higher temporal resolution of every 2 hours at 5 dpf using two-photon imaging (Fig 4a, Supplementary Movie 14,15). We traced every *nr1d1*:VNP-positive cell at different time of the day (Fig. 4b, Supplementary Table 3). Fig. 4c showed the trace plots of individual cells of two zebrafish throughout the day. Using cosine fitting P<0.05 and relative oscillating amplitude>0.05 as cutoff, we identified 84 cells showed circadian oscillations in a total of 117 *nr1d1*:VNP-positive cells in the pineal gland. Their circadian phases were distributed around ZT11 within a narrow range (Fig. 4d) with Kuramoto order parameter R=0.86 indicating the phase coherence of single-cell circadian oscillators [21]. Clustering analysis of single-cell VNP traces identified two distinct clusters of cells that can be distinguished by their baseline fluorescence intensity (Fig 4e, f, g). Such difference in baseline level of expression at 5dpf can be explained by the cell-type specific developmental trajectories that we observed above. We also observed that the cluster with higher baseline level (cluster 1) also showed lower relative amplitude of oscillation (Fig 4h). Similar negative correlation between mean transcriptional activity and relative amplitude of circadian oscillations has been reported in mouse [22]. However, there is no significant difference in circadian phase between two clusters (Supplementary Fig. 3). In short, circadian clocks in pineal photoreceptor cells are oscillating synchronously under LD in spite of large difference in baseline level.

**Fig. 4.**
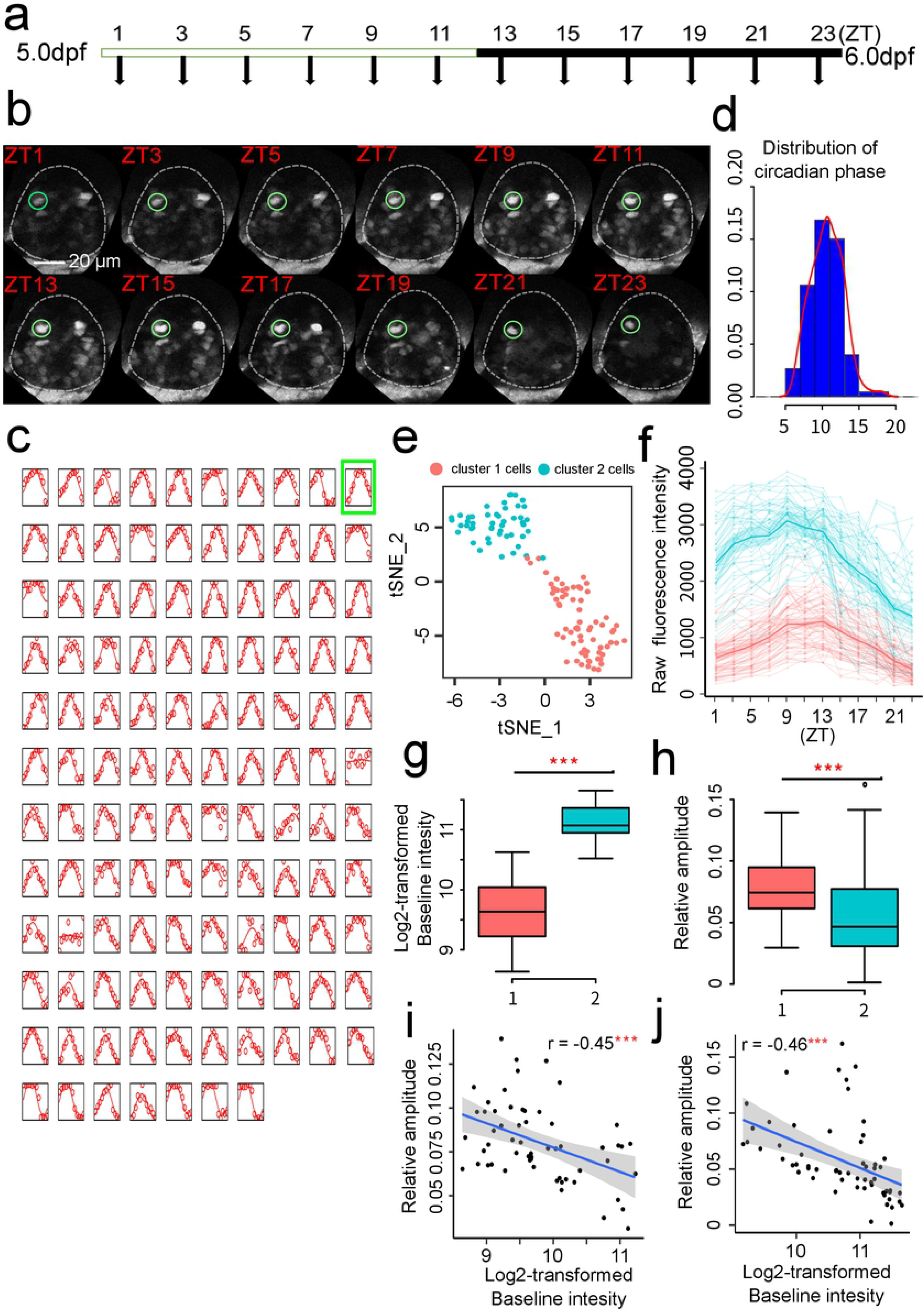
Circadian dynamics of pineal gland at higher temporal resolution. (a) Experimental design to examine the developmental dynamics of *nr1d1*:VNP expression at higher temporal resolution. (b) Fluorescence images illustrate the tracing of one *nr1d1*:VNP example cell every 2 hours. (c) Single-cell tracing results of all 117 *nr1d1*:VNP-positive cells in two zebrafish pineal glands (2 fish). The empty dots represent the original fluorescence signals, while the solid line represents the smoothed curve fitted by the cosine functions. The example cell in (b) is highlighted by a green square. (d) Circadian phase distributions of the 117 *nr1d1*:VNP-positive cells. (e) t-SNE visualization of the clustering result of the 117 *nr1d1*:VNP-positive cells. (f) Raw fluorescence intensity traces of the two types of *nr1d1*:VNP-positive cells in (e). (g & h) The Comparison of the baseline expression (g) and relative circadian amplitude (h) between the two types of cells. The colors of the boxes corresponds to (e). Two-tailed Student’s t-test was applied to calculate the levels of significance between the two types of cells. *** represents P<0.001. (i & j) Scatter plot demonstrated the relationship between baseline intensity and relative oscillation amplitude for fish1 (i) and fish2 (j), respectively. *** represents Pearson’s correlation p-value <0.001.

### Light-Dark cycle is essential for the onset of cellular circadian oscillation

It is known that LD cycle is required for the development of circadian clock in zebrafish [9]. However, it is unclear whether the role of LD cycle in clock ontogeny is a synchronization of individual oscillators or an initiation of oscillation. To address this question, we imaged *nr1d1*:VNP signals in zebrafish larvae raised under the constant dark (DD) from 3.5 dpf to 6.5 dpf using two-photon imaging (Fig. 5a, Supplementary Movie 10-13, Supplementary Table 6). Examination of the expression patterns of all *nr1d1*:VNP-positive cells from four DD fish showed that although the expression of *nr1d1*:VNP still increased during development, the circadian expression had been absent in DD cells (Fig. 5b). Indeed, regression analysis showed that the oscillating coefficients of majority of DD cells are not significantly different from zero (Fig. 5c). However, DD cells have even higher developmental coefficients than LD cells (Fig. 5d). In comparison, when we imaged the fish transferred into DD condition after six LD cycles (LD_DD cells) (Fig 5e, Supplementary Table 7-8, Supplementary Movie 19-24), we observed that LD_DD cells were still oscillating into two days of darkness (Fig 5f) and the amplitude of circadian oscillation showed no significant difference comparing to the fish kept under LD condition (LD_LD cells) (Fig 5g). To further examine the circadian oscillation in dark-raised fish, we imaged *nr1d1*:VNP signals in the pineal gland every 2 hours at 5 dpf (Fig 5h, Supplementary Movie 16-18, Supplementary Table 4). We found that DD cells showed much lower oscillating amplitudes than those in LD cells (Fig. 4i, j). Only 1 out of 142 DD cells showed significant oscillation (p-value<0.05 and relative amplitude>0.1, Fig 5k). Therefore, our result suggests that the cellular circadian oscillations in DD fish were not initiated rather than not synchronized during development.

**Fig. 5.**
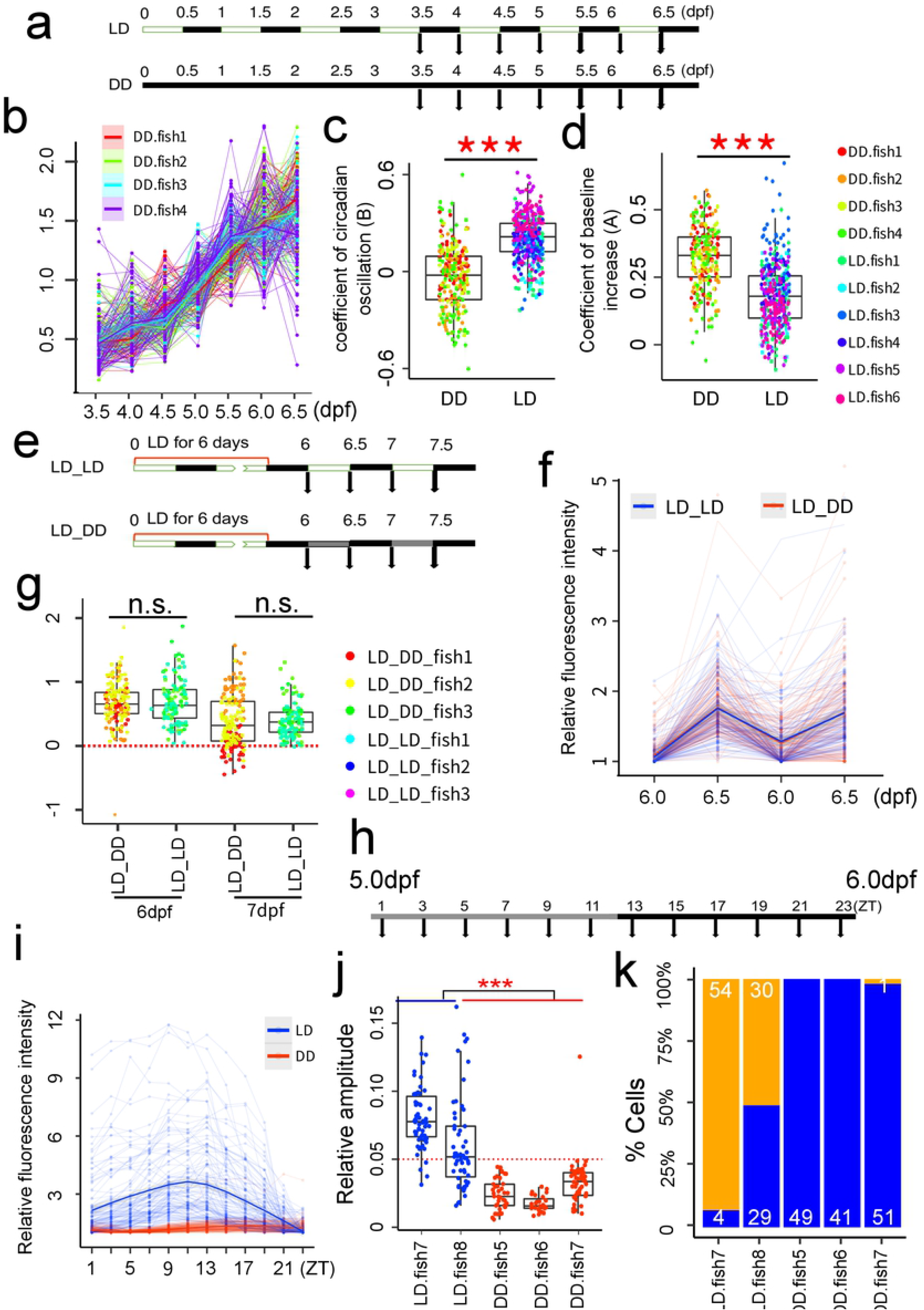
Light-Dark cycle is essential to synchronous onset of *nr1d1* oscillation. (a) Experimental design to investigate the effect of light on the onset of circadian clock development. (b) Expression patterns of all cells under DD condition during development (four fish). Each thin line represents one cell. Each thick line represents the loess-smoothed curve of all cells in each individual fish. The shaded area shows the 95% confidence level of the smoothed curve. (c) Comparison of oscillating coefficient (B) between LD and DD cells. Two-tailed Student’s t-test was applied to calculate the levels of significance between the two types of cells. *** represents P<0.001. (d) Comparison of developmental coefficient (A) between LD and DD cells. Two-tailed Student’s t-test was applied to calculate the levels of significance between the two types of cells. *** represents P<0.001. (e) Experimental design to examine the single-cell circadian clocks under DD after transferred from LD (LD_DD). (f) Expression patterns of LD_LD (three fish) and LD_DD cells (three fish) from 6.0 to 7.5dpf. Each thin line represents one cell and each dot represents raw fluorescence intensity. Thick lines represent the loess-smoothed curves for all the LD_DD cells in red and LD_LD cells in blue respectively. The shaded areas show the 95% confidence level of the smooth curve. (g) Log2-transformed dusk-dawn ratios of VNP fluorescence intensities of all the cells in (f). Each data point represents one cell and the colors correspond to different fish. The red dash line represents y=0. n.s. represents no significant difference in dusk-dawn ratios between different time points (two-tailed ANOVA test). (h) Experimental design to examine the expression pattern of DD cells at higher temporal resolution. (i) Expression patterns of LD (two fish) and DD cells (three fish) across one day at 2-h resolution. Each thin line represents one cell and each dot represents normalized fluorescence intensity by dividing minimal value of each cell. Thick lines represent the loess-smoothed curves for all the LD in blue and DD cells in red respectively. The shaded areas show the 95% confidence level of the smooth curve. (j) Comparison of the relative amplitudes of LD and DD cells in (j). The red dash line represents y=0.05. (k) Percentages of oscillating cells (cosine p-value<0.05 and relative amplitude>0.05) in each LD and DD fish. The orange bars represent the percentage of oscillating cells while the blue bars represent the percentage of non-oscillating cells.

## Discussion

In our study, we constructed a zebrafish *nr1d1*:VNP fluorescence reporter system and for the first time we can monitor the circadian gene expression at the single-cell resolution in zebrafish larvae. Using this reporter fish, we revealed the interplays among circadian clock, cell-type specific development, and light-dark cycle. The mouse version of *nr1d1*:VNP was originally developed by Nagoshi et al. in NIH 3T3 cells. VNP was chosen for its high fluorescence intensity and high folding efficiency [23]. With this reporter, they were able to show that each NIH 3T3 fibroblast cell has a self-sustained circadian oscillator that can be entrained by serum shock. Also using this reporter, two independent studies have demonstrated the tight coupling between circadian rhythm and cell cycle in proliferating NIH 3T3 cells [24,25]. As FUCCI lines to label different stages of cell-cycle are available in zebrafish [26], one can cross *nr1d1*:VNP fish with cell-cycle reporter lines to examine the relationship between circadian clock and cell cycle *in vivo*. By crossing fish lines containing cell-type specific fluorescent markers with our *nr1d1*:VNP fish, one can conduct cell-type specific imaging of circadian rhythm as we have done for rod cells in pineal gland. Larval zebrafish has also been used in drug screening for chemical compounds affecting circadian rhythm and sleep [27]. Our Nr1d1:VNP fish can be used to investigate the effect of selected drug targets on the synchronization of single-cell oscillators *in vivo*. With advanced microfluidic systems to capture and recapture larval zebrafish, one can conduct more intense imaging on the same fish to achieve higher temporal resolution while minimizing the perturbations to the fish [28]. Using newly developed movable platform to track zebrafish larvae and novel volume imaging system, live imaging of cellular circadian rhythm on free-moving zebrafish will become possible [29]. In summary, our single-cell circadian reporter zebrafish line will have broad applications in circadian research.

It was believed that physiological and behavioral rhythms in zebrafish appear during the early stages of zebrafish development [8]. In our study, the expression of *nr1d1* appeared gradually in development and step-wise in different brain regions. As the major component of the second negative feedback loop in circadian clock, *nr1d1* is directly regulated by BMAL1 and represses the expression of *bmal1* [1]. The onset of *nr1d1* expression may represent the maturation of a functional circadian clock during development. Our single-cell study revealed the developmental trajectory of *nr1d1* expression varies greatly among individual cells and the onset of circadian rhythm is dependent on early light exposure. Ziv et al. has shown that *per2* expression can be induced by light and is necessary for light-dependent onset of circadian oscillation of *aanat2* in the pineal gland [30]. In addition, *otx5*, a transcription factor constantly expressed in circadian cycle, has been shown to regulate the expression of *nr1d1* in pineal gland [31]. It is likely *otx5* is responsible for cell-type specific developmental components of *nr1d1* expression. Nagoshi et al. have suggested that the robust expression of *nr1d1*:VNP in mouse NIH 3T3 cells is due to high transcriptional activity of *Nr1d1* promoter in these cells as *Nr1d1* is more highly expressed in tissues such as liver compared to other core clock genes such as *Bmal1* [23]. In our case, the presence of *otx5* and *crx* binding sites in *nr1d1*:VNP promoter may have given rise to its robust expression in pineal photoreceptors and cells in other brain regions. Therefore, one can replace *otx5* and *crx* binding sites by those of other cell-type specific transcription factors in *nr1d1*:VNP to report the circadian rhythm in cell types of interest. This will enable single-cell imaging of circadian clock in a broad range of cell types in live zebrafish.

In Ziv et al.’s study, they proposed that the expression of *aanat2* is asynchronous in individual pineal cells under constant darkness [30]. By imaging *per1-luc* transgenic fish, Dekens et al. also suggested that cells in the embryos kept under constant environmental conditions undergo asynchronous oscillations [32]. Carr and Whitmore showed that individual cells of zebrafish cell line kept under long-term darkness became unsynchronized [10]. To our surprise, we did not observe asynchronous oscillations of *nr1d1*:VNP among individual cells in our reporter fish raised under constant darkness suggesting that cellular oscillators were not initiated without environmental stimuli. Our study provides direct evidence that cellular oscillators are initiated by light exposure during *in vivo* clock development demonstrating the utility of single-cell circadian zebrafish reporter.

## Methods

### Construction of the *nr1d1*:VNP (Venus-NLS-PEST) Expression Vector

The zebrafish *nr1d1*:VNP reporter plasmid was constructed as follows. The mouse *Nr1d1*:VNP reporter plasmid was generously provided by Emi Nagoshi [23] (Department of Genetics and Evolution, Sciences III, University of Geneva). The plasmid carries an intact VNP sequence. VNP (*V*enus-*N*LS-*P*EST) is a nuclear fluorescent protein with a high folding efficiency and short half-life [24]. The zebrafish *Nr1d1*:VNP plasmid was obtained by homologous recombination with four cassettes. Targeting cassette was generated by four-step PCR amplification. The first PCR step was performed to amplify the tol2-polyA-Ampicillin gene with an additional 3′ sequence that is homologous to the 5 ′ end region of zebrafish *nr1d1* promoter (template: 394-tcf-lef-TuboGFPdest1, forward primer: 5′ - GCTTCTGCTAGGATCAATGTGTAG-3′, reverse primer: 5′ - CCTATAGTGAGTCGTATTACCAACTT-3′). The second and third PCR steps were performed to amplify the promoter of zebrafish *nr1d1* gene (6.2k) (template sequences from zebrafish genome: second PCR forward primer: 5′ - TACGACTCACTATAGGTTTTCCACCCGCTGGGCTGCCTCTCACGTG-3′, reverse primer: 5′-TATTCCTTGCTTCCGTCTGTAGTGCCCAAT-3′; third PCR forward primer: 5′-GGCAGACATCACGGGTTAAGACACAGTGTT-3′; reverse primer: 5′ - CAGCAGGCTGAAGTTAGTAGCTCCGCTTCCTCCCGGAGGGGAT-GGTGGGAATGATGATCC-3′). The fourth PCR step was performed to amplify the cassette of VNP (template sequences from mouse *Nr1d1*:VNP reporter plasmid forward primer: 5′-ACTAACTTCAGCCTGCTGATGGTGAGCAAGGGCGAGGA-3′, reverse primer: 5′-CTACACATTGATCCTAGCAGGAGCACAG-3′). The 3′ arm of the first PCR was homologous to the 5′ arm of the second PCR, the 3′ arm of the second PCR was homologous to the 5′ arm of the third PCR, the 3′ arm of the third PCR was homologous to the 5′ arm of the fourth PCR, and the 3′ arm of the fourth PCR was homologous to the 5′ arm of the first PCR. The vector construct was verified by sequencing. Fig. 1a shows the schematic of *nr1d1*:VNP construct design. The structure of 6.2k-bp *nr1d1* promoter containing multiple E-box, RRE, and pineal specific elements was also described.

### Generation of transgenic Tg(*nr1d1*:VNP) zebrafish reporter line

Generation of transgenic fish using the Tol2 system was carried out according to the published protocol [33]. Constructs were injected together with capped RNA encoding transposase (10 ng/μl each of DNA and RNA) into fertilized eggs at the one-cell stage. The injected fish (F0 generation) were raised and screened for integration of the transgene into the germline. Isolation of transgene-positive progeny (F1) was carried out using EGFP imaging using fluorescence stereomicroscpe (Olymous, SZX16). For *in vivo* imaging, *Tg(nr1d1*:VNP) zebrafish was crossed with a pigmentation mutant strain (*casper* mutant), which is transparent in the whole brain except the eyes.

### Generation of transgenic Tg(lws2:nfsB-mCherry) and Tg(xops:nfsB-mCherry) zebrafish reporter line

The 1.77 kb lws2 and 1.38 kb xops promoter were amplified by PCR. Specific primers for the lws2 promoter: 5’-ggccagatgggccctGTTGTGCACCAGATCTGAGT-3’ and 5’-TGGTCCAGCCTGCTTTTTGGAAACCCTGAAGATCA-3’; Specific primers for the xops promoter: 5’-ggccagatgggccctGCGGCCGCAGATCTTTATACAT-3’ and 5’-TGGTCCAGCCTGCTTCCTCGAGATCCCTAGAAGCCT-3’; The products were subcloned into pTol-uas:nfsB-mCherry plasmid to substitute the uas promoter by using homologous recombination kit (ClonExpress MultiS One Step Cloning Kit, C113; Vazyme, China). The pTol-lws2:nfsB-mCherry plasmid and pTol-xops:nfsB-mCherry were co-injected into AB embryos with Tol2 transposase mRNA at one-cell stage. The mCherry signal was screened to identify F1.

### Zebrafish husbandry

Adult zebrafish (NCBI taxonomy ID: 7955) were raised and maintained in fully automated zebrafish housing systems (Aquazone, Israel; temperature 28 °C, pH 7.0, conductivity 300 mS) under 14h/10h LD cycles, and fed with paramecium twice a day. Larvae were fed twice a day starting from 5dpf. For the experiments under normal LD condition, embryos were produced by natural spawning in the morning and raised in egg-water containing methylene blue (0.3 ppm) in a light-controlled incubator under 12h/12h LD cycles at 28 °C. ZT0 is defined as the time when the lights are turned on (9 A.M.). For the experiments under DD condition, embryos were collected in 30 minutes after birth and put into a black box in a dark incubator at 28 °C. All the experimental protocols were approved by the Animal Use Committee of Institute of Neuroscience, Chinese Academy of Sciences.

We use Tg(*nr1d1*:VNP), Tg(*aanat2*:mRFP), Tg(*lws2*:nfsB-mCherry), Tg(*xops*:nfsB-mCherry), and Tg(*her4*:DsRed) zebrafish lines maintained on a AB background. Sex includes male and female. We used the larva fish aged from 3.5 dpf to 7.5 dpf.

### In vivo imaging

Fish were randomly selected to be imaged under LD or DD. We were not blinded to LD and DD fish group allocation. For time-lapse imaging in every two hours lasting for one day starting at ZT0, 5.0dpf zebrafish larvae were anesthetized with MS-222 (sigma-Aldrich) (0.01-0.02%) and mounted in low melting-point agarose (1.33%). The fish were maintained in a chamber with circulating egg-water without anesthesia at temperature of 28 ± 0.5 °C. For imaging from 3.5dpf to 6.5dpf every 12 hours at ZT0 and ZT12, the zebrafish larvae were anesthetized at each time point with MS-222 (sigma-Aldrich) (0.01-0.02%) and imbedded in low melting-point agarose (0.8–1.0%) before imaging. 3-4 fish were imaged in each imaging session that lasts for half to one hour. Fish were released immediately after the end of each imaging session and in a free-moving state between image sessions. All fish were imaged from dorsal view using two-photon microscope (Olympus) as described previously [34]. Imaging was performed using a 25× (numerical aperture (NA)=1.05) water-immersion objective (Olympus). Excitation was provided by a Ti:Sapphire femtosecond pulsed laser system (Coherent) tuned to 900 nm, which allowed efficient simultaneous excitation of Venus fluorescent protein. Laser power was set to 1.6% for pineal imaging and 2.0% for whole brain imaging. We use two-photon excitation microscopy for two reasons as suggested by Carvalho and Heisenberg [35]. First, the near-infrared wavelength has undetectable effects on fish physiology and behavior. Second, as the two-photon excitation is only achieved near the focal plane, it minimizes photo-bleaching and photo-toxicity. The sample size estimate is based on our previous studies. Two fish under light-dark (LD) condition were imaged for the whole brain imaging. Two fish under LD condition and three fish under dark-dark(DD) condition were imaged for one day every 2 hours. Six fish under LD condition and four fish under DD condition were imaged from 3.5 dpf to 6.5 dpf every 12 hours. Three fish under LD_LD condition and Three fish under LD_DD condition were imaged from 6.0 dpf to 7.5 dpf every 12 hours. Six rod fish were imaged from 3.5 dpf to 6.5 dpf every 12 hours.

### Single cell RNA-seq and data analysis

6dpf larval heads were dissected on dissection medium (DMEF/F12 with 2% 100X penicillin-streptomycin) and pineal region were enriched by pipetting the pineal into the tube. Dissociation of the brain cells following the protocol from Miguel A. Lopez-Ramirez et al. [36]. Briefly, Add 300ul papain solution to the dissected tissue at 37 in water heater for 15 minute, pipetting during digest. Then stop the digestion by adding 1.2ml of washing solution. Washing the cells twice using washing solution. In the end, sterilize the cells using 40um pore size filter. Stain cells using trypan blue and count the living cells using a hemocytometer. All the reagents and solutions used were following the materials provided by Miguel A. Lopez-Ramirez et al. [36]. The resulting single cell suspension was promptly loaded on the 10X Chromium system using Chromium Single Cell 3’ Reagents v2.

For data analysis, raw sequencing data was converted to matrices of expression counts using the cellranger software provided by 10X Chromium. Reads were aligned to a zebrafish reference transcriptome (ENSEMBL Zv11, release 95). The gene expression matrix were then loaded into Seurat package [37] in R for the following analysis. Cells with less than 500 genes or percentage of mitochondiral genes>0.02 were excluded from the following analysis. The left cells were clustered using the first 50 PCs and with resolution=1, 26 clusters were identified. They were manually annotated by comparing the marker genes with those from whole-brain single-cell RNA-seq data in adult zebrafish [19].

### Image analysis

The image data were first converted to ‘nrrd’ format using a customer Fiji [38] macro code for downstream analysis. Time series of 3D images in ‘nrrd’ format for each fish were then aligned by CMTK toolkit (parameters: -awr 01 −l fa -g -T 8 -X 10 -C 1 -G 20 -R 2 -E 1 -A ’--accuracy 5 --dofs 12’ -W ’--accuracy 5’ for the alignment of pineal gland; -ar 01 −l a -A ’--exploration 50 --accuracy 5 --dofs 12’ for the alignment of whole brain) to facilitate the single cell tracing. TrackMate [39], an open source Fiji [38] plugin, was applied to perform the single cell nucleus identification and tracing on 3D images over time. Briefly, Laplacian of Gaussian (LoG) filter with a sigma suited to the estimated blob diameter (5.5 micron) was applied to detect the single cell nuclei in each 3D image. Then HyperStack Displayer was applied to visualize the identified spots, which allows manual editing afterwards. A nearest-neighbor algorithm (parameter: maximal linking distance 5.5 micron) was applied to trace each single cell. After the automatic processing, the identification and tracing of each cell were manually curated using the manual tracking tools of TrackMate [39]. Combining the automated and manual tracking approaches can significantly increase the detection accuracy of single cells. In the end, the cell position and mean image intensities were exported for the following statistical analysis. For the whole brain data, the pineal gland, the optic tectum, and the cerebellum were first manually extracted, then single cells were traced using manual tracking tools of TrackMate [39], and mean intensities of each brain region was used for the following analysis. The cell position and mean intensities for all the fish analyzed were listed in Supplemental Table 2-9.

### 3D reconstruction of zebrafish pineal gland

To systematically analyze the 3D distribution of the *nr1d1*:VNP-positive cells, a common 3D pineal gland structure was reconstructed by averaging six zebrafish pineal glands at 5.5dpf. First, we manually reconstructed the 3D pineal structure of six individual 5.5dpf zebrafish pineal glands taking advantage of the clear pineal gland boundary in two-photon imaging. Secondly, all 3D pineal gland structures from different fish were aligned using rigid transformation using CMTK (parameters: -ar 01 −l a -A ’--accuracy 5 --dofs 12’). Thirdly, the signals from aligned 3D pineal glands were added and smoothed by Gaussian blur (Sigma = 30) using Fiji package [38]. In the end, a common 3D pineal gland structure was obtained by implementing Li’s minimum cross entropy thresholding method using Fiji package [38].

The cells of each pineal gland were then registered to the common 3D pineal gland using the same transform matrix for the alignment of the 3D pineal gland structure. The ‘ks’ package in R (https://www.R-project.org/) was applied to calculate the cell density distribution. The common zebrafish pineal structure and *nr1d1*:VNP cell density distribution were visualized by NeuTube software [40] (Fig. 2d).

### Statistical analysis

All the statistical analysis was performed using the computing environment R (https://www.R-project.org/). For the time-lapse imaging data from 5.0 dpf to 6.0 dpf every 2 hours, the circadian phase and amplitude of each cell were calculated by fitting the mean intensity of each cell to cosine functions with 24 hours’ period and shifting phases as described previously [41]. The fluorescence intensities were log2-tranformed before fitting. Relative amplitude were defined as absolute amplitudes divided by means. R package ‘circular’ was applied to calculate the average circadian peak (mean.circular) and Kuramoto’s order parameter R (Rho.circular). For the imaging data of developmental time course from 3.5dpf to 6.5dpf, the fluorescence intensities were first normalized by dividing the mean intensity. A regression model combining a stepwise function denoting the circadian effect and a linear function denoting the developmental trend were used to fit the data. Namely, F(*x*) = Ax+ B*(−1)^(*x*+1)^ + C, A denotes the slope of developmental trend, B denotes the amplitude of circadian oscillation, and C denotes the baseline level of fluorescence intensity. Nonlinear least squares (nls) were applied to determine these three parameters and the level of significance of each coefficient. R package ‘Seurat’ was applied to perform t-SNE analysis and visualization of clusterings. To classify LD cells imaged every 2 hours, all the twelve principle components were used and resolution was set to 0.4. calculate the levels of significance in differences between the two types of cells (Fig. 3e and 3f) and between the LD and the DD cells (Fig. 3g, h and 4g, h).

### Real-time qPCR

For each time point, A total of 15-20 fish were separated into 2 tubes equally. Here the sample size estimate is based on our previous studies. Trizol (Invitrogen) was immediately added to each tube. Then total RNA was isolated from zebrafish larvae by Trizol (Invitrogen) according to manufacturer’s protocol. Total RNA quantities were measured by a Nanodrop spectrometer (Nanodrop 2000). 500 ng of total RNA was reverse transcribed using HiScript II Q RT SuperMix for qPCR (+gDNA wiper) (vazyme) according to manufacturer’s protocol. 2 μ l of RT product (1:10 diluted) and 10μl of SYBR Green Master Mix (vazyme) were used in qPCR on a ABI StepOne Plus System (Thermo Fisher) according to manufacturer’s protocol. The specificity of PCR was checked by melting curve analysis. In every qPCR assay, *eef1a1* was used as the control gene for any significant bias of starting materials across samples. Primers used in this study were listed in Supplementary Table 10.

## Acknowledgement

This work was supported by National Natural Science Foundation for Young Scientists of China Grant (No. 31701029) and Natural Science Foundation of Shanghai Grant (16ZR1448800) and to H.-F.W; National Natural Science Foundation of China Grant (No. 31571209 to J.Y and No. 81401279 to Z.-Y. Y). We thank Prof. Emi Nagoshi (Department of Genetics and Evolution, Sciences III, University of Geneva, Switzerland) for the mouse *Nr1d1*:VNP reporter plasmid and Prof. Yoav Gothilf (Department of Neurobiology, The George S. Wise Faculty of Life Sciences and The Sagol School of Neuroscience, Tel-Aviv University, Israel) for *aanat2*:mRFP transgenic zebrafish.

## Author contributions

H.W. performed part of the experiments and did all the analyses. Z.Y., X.L., D. H., and S. Y. performed most of the experiments. J.Y., J.H., and Y.L. conceived the study and experimental design. J.Y., J.H., and H.W. wrote the manuscript.

## Code availability

All codes for imaging analysis and statistical analysis are available upon request.

## Data availability

Sigle cell RNA-seq data has been deposited on GEO (GSE134288). All the compressed original images are provided as Supplementary Movies. All the original tiff files are available upon request. All the single cell tracing results are provided as Supplementary Tables.

